# Dopamine D1 receptor-expressing neurons activity is essential for locomotor and sensitizing effects of a single injection of cocaine

**DOI:** 10.1101/2020.12.19.423602

**Authors:** Yukari Nakamura, Sophie Longueville, Akinori Nishi, Denis Hervé, Jean-Antoine Girault, Yuki Nakamura

## Abstract

D1 dopamine receptors play an important role in the effects of cocaine. Here we investigated the role of neurons which express these receptors (D1-neurons) in the acute locomotor effects of cocaine and the locomotor sensitization observed after a second injection of this drug. We inhibited D1-neurons using double transgenic mice conditionally expressing the inhibitory Gi-coupled *designer receptor exclusively activated by designer drugs* (Gi-DREADD) in D1-neurons. Chemogenetic inhibition of D1-neurons by a low dose of clozapine (0.1 mg/kg) decreased the induction of *Fos* in striatal neurons. It diminished the basal locomotor activity and acute hyper-locomotion induced by cocaine (20 mg/kg). Clozapine 0.1 mg/kg had no effect by itself and did not alter cocaine effects in non-transgenic mice. Inhibition of D1-neurons during the first cocaine administration reduced the sensitization of the locomotor response in response to a second cocaine administration ten days later. At day 11, inhibition of D1-neurons by clozapine stimulation of Gi-DREADD, prevented the expression of the sensitized locomotor response, whereas at day 12, in the absence of clozapine and D1-neurons inhibition, all mice displayed a sensitized response to cocaine. These results show that chemogenetic inhibition of D1-neurons decreases spontaneous and cocaine-induced locomotor activity. It blunts the induction and prevents the expression of sensitization in a two-injection protocol of sensitization but does not alter established sensitization. Our study further supports the central role of D1-neurons in mediating the acute locomotor effects of cocaine and its sensitization.

## Introduction

Psychostimulants such as cocaine increase locomotor activity and repeated cocaine exposure progressively increases the locomotor responses due to sensitization processes (review in Steketee and Kalivas, 2011). Locomotor sensitization to repeated drug administration is a complex adaptation that involves several neuronal populations and is correlated with aspects of addiction including the excessive pursuit and self-administration of psychostimulant drugs (Vezina, 2004). However, multiple drug exposures can promote the parallel development of tolerance that could interfere with behavioral sensitization. Remarkably, locomotor sensitization is visible even following a single cocaine administration, as evidenced by a two-injection protocol of sensitization (TIPS, Jackson and Nutt, 1993; Valjent et al., 2010). TIPS is a simple protocol to explore the mechanisms by which drugs of abuse exert long-lasting effects. It has the advantage to allow simple manipulations during the unique initial injection and test their consequences on subsequent sensitization.

Dopamine D1 receptors (Drd1) play a major role in the rewarding properties of drugs of abuse including cocaine (Self et al., 1996). Cocaine treatment stimulates the activity of D1-neurons in the striatum as evidenced by the rapid Drd1-dependent increase in intracellular calcium selectively observed in these neurons in vivo (Luo et al., 2011). SCH23390, a Drd1 antagonist that also acts on other receptors, prevents locomotor sensitization (McCreary and Marsden, 1993). *Drd1* knockout mice lack locomotor response to psychostimulants (Drago et al., 1996; Miner et al., 1995) and display altered locomotor sensitization after repeated cocaine administration (Karlsson et al., 2008). With the TIPS protocol, sensitization (20 mg/kg) was absent in homozygous *Drd1* knockout mice, and reduced in heterozygous mice (Valjent et al., 2010). However there is a limitation in the interpretation of knockout mice experiments, because the lack of *Drd1* gene affects the expression of other genes (Xu et al., 1994) and developmental alterations cannot be excluded. In addition to the role of Drd1, conditional deletion of the NR1 subunit of the glutamate NMDA receptor in D1-neurons impaired amphetamine sensitization (Beutler et al., 2011). These experiments provide evidence about the role of receptors but do not explore the contribution of specific neuronal populations. Several cell populations are implicated in behavioral sensitization. Pharmacological evidence shows that local administration of amphetamine into the VTA induces Drd1-dependent locomotor sensitization (Vezina, 1996). Unexpectedly, locomotor sensitization to amphetamine was prevented in mice in which dorsomedial striatum Drd2 neurons were ablated but not in those with lesion of D1-neurons in the dorsal striatum (Durieux et al., 2012). Metamphetamine sensitization was delayed but not prevented in mice in which D1-neurons were ablated in the nucleus accumbens (NAc) shell (Kai et al., 2015). Cocaine locomotor sensitization was markedly decreased in mice in which neurotransmission was selectively impaired in NAc D1-neurons (Hikida et al., 2010) and optogenetic inhibition of NAc D1-neurons also attenuated this response (Chandra et al., 2013). Potentiation of corticostriatal synapses onto NAc D1-neurons is a key component of locomotor sensitization in response to a single injection of cocaine (Pascoli et al., 2011). However recent work reveals that other striatal neurons, cholinergic interneurons, also play a role in responses to cocaine, including sensitization (Lewis et al., 2020).

In the current study we asked two simple questions: Does inhibition of D1-neurons alter i) the acute locomotor response to cocaine and ii) the induction or expression of sensitization in the TIPS protocol? To address these questions we used chemogenetics with designer receptor exclusively activated by designer drug (DREADDs) (Roth BL et al., 2016) and expressed hM4Di, an inhibitory DREADD, selectively in D1-neurons. Since recent work showed that DREADD stimulation with low doses of clozapine (CLZ) is faster and at least as efficient as clozapine-N-oxide (CNO), the commonly used ligand (Gomez et al., 2017), an observation confirmed in our laboratory (Nakamura et al., 2020), we used CLZ to activate hM4Di and inhibit D1-neurons.

## Materials and Methods

### Animals and treatments

We used heterozygous bacterial artificial chromosome (BAC) *Drd1*-Cre transgenic (Drd1-Cre) mice expressing Cre recombinase under the control of the dopamine D1 receptor gene (*Drd1*) promoter (GENSAT project, EY262 line), bred onto a C57BL/6J genetic background. Gi-DREADD (B6N.129-*Gt(ROSA)26Sor*^*tm1(CAG-CHRM4*,-mCitrine)Ute*^/J, a.k.a. R26-LSL-hM4Di-DREADDpta-mCitrine or R26-hM4Di/mCitrine) mice were purchased from the Jackson Laboratory (stock #026219). We crossed Drd1-Cre mice with Gi-DREADD mice. A subset of their offspring carried both *Drd1*-Cre and R26-hM4Di/mCitrine alleles, allowing effective Cre-dependent removal of the stop cassette and transient inhibition of Gi-DREADD-expressing neurons upon CLZ treatment (**Fig. 1A**). Male and female offspring (10-21-week-old at the beginning of the experiments) were used for the study. Genotypes were determined by polymerase chain reaction (PCR) on mouse tail DNA samples. Cocaine-HCl (Sigma, France) was dissolved in 9 g.L^−1^ NaCl solution (saline). CLZ (Sigma-Aldrich) was first dissolved in DMSO, and then diluted in saline (final DMSO concentration, 0.1 mL.L^−1^). Cocaine-HCl (20 mg.kg^−1^) and CLZ (0.1 mg.kg^−1^) were administered by intraperitoneal injections in a volume of 10 mL.kg^−1^ body weight. Mice were housed under a 12-hour light/12-hour dark cycle with free access to food and water. All animal procedures used in the present study were approved by the *Ministère de l’Education Nationale de l’Enseignement Supérieur de la Recherche* (France, project APAFIS#8196-201704201224850 v1. All methods in this study were performed in accordance with the relevant guidelines and regulations.

**Figure 1.**
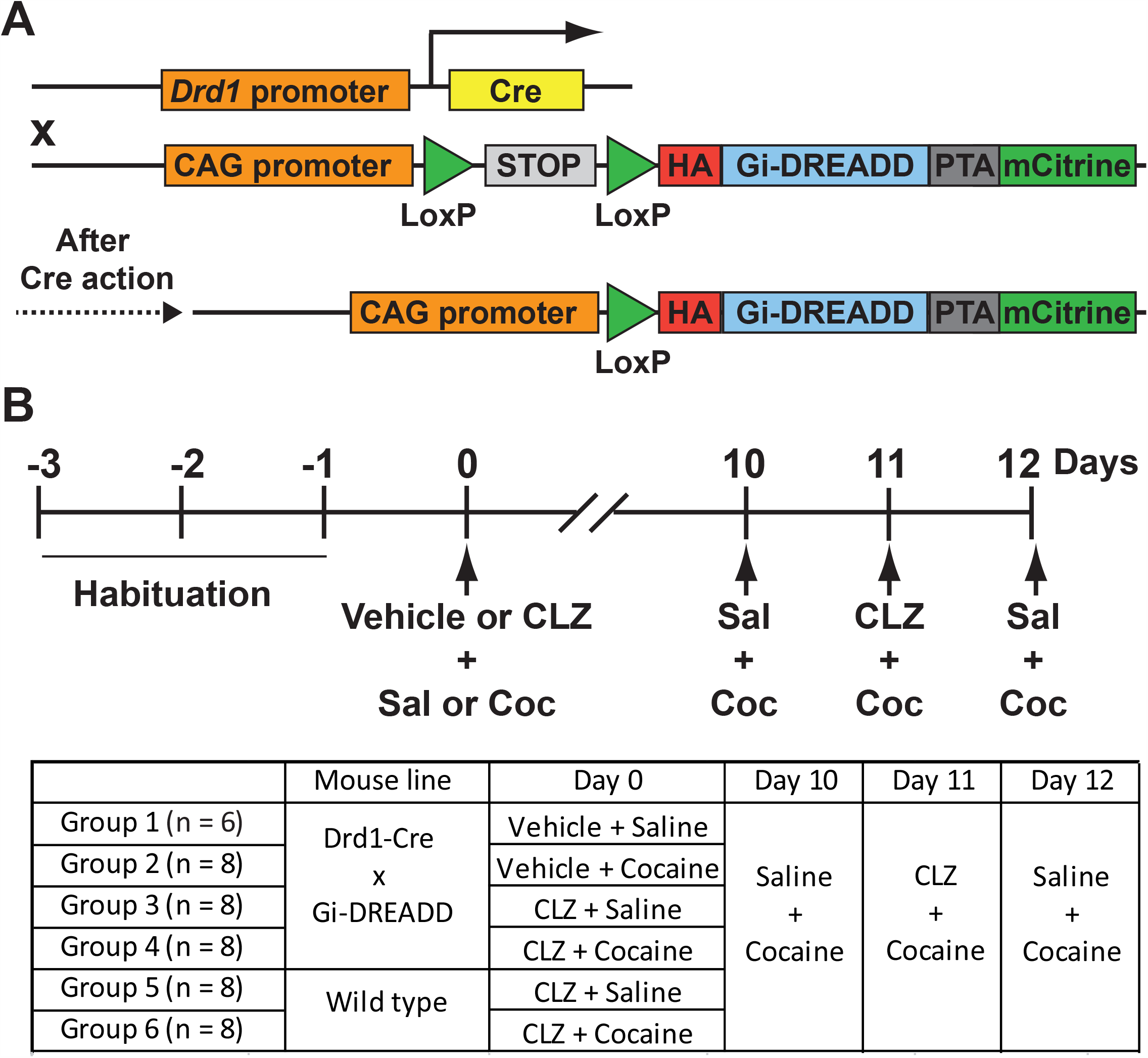
Schematic experimental design. **A)** Gi-DREADD (hM4Di) expression in Drd1-expressing neurons (D1-neurons) of double transgenic Drd1-Cre x Gi-DREADD mice. Following Cre-mediated removal of an upstream floxed-STOP cassette, the HA-tagged Gi-DREADD and mCitrine are expressed in D1-neurons. HA, hemagglutinin epitope tag, PTA, porcine teschovirus cleavage site. **B**) Schematic schedule of behavioral experiments and groups of mice used in the study. Sal, saline, Coc, cocaine (20 mg/kg), CLZ, clozapine (0.1 mg/kg).

### Behavioral analysis

Locomotor activity was measured in a circular corridor with four infrared beams placed at every 90° angles (Imetronic, Pessac, France) in a low luminosity environment (locomotor activity box, LA box). Counts were incremented by consecutive interruptions of two adjacent beams. Locomotor sensitization induced by a single cocaine injection was studied as described previously (Valjent et al., 2010). All mice were habituated to handling, test apparatus, and procedures for 3 consecutive days (Days −3 to −1, **Fig. 1B**) before the actual experiment. In this habituation period, mice received every day a first saline injection and were placed in the LA box for 30 min. Then, they received a second saline injection and were put back in the box for 1 h. On experimental days (Day 0, Day 10, Day 11, and Day 12), the animals were handled in the same manner, except that they were injected with the solutions indicated in **Fig. 1B** at 0 min and 30 min, respectively.

### Tissue preparation and immunofluorescence

Drd1-Cre x Gi-DREADD mice were injected with vehicle or CLZ and placed in the LA box for 30 min. Then, they received a cocaine injection and were placed back in the box for 90 min, after which they were injected with a lethal dose of pentobarbital (500 mg.kg^−1^, i.p., Sanofi-Aventis, France) and perfused transcardially with 40 g.L^−1^ paraformaldehyde in 0.1 M sodium phosphate buffer (**Fig. 2A**). Brains were post-fixed overnight at 4°C, cut into free-floating sections (30 μm) with a vibrating microtome (Leica) and kept at −20°C in a solution containing 300 mL.L^−1^ ethylene glycol, 300 mL.L^−1^ glycerol in 0.1 M phosphate buffer. Immunolabeling procedures were as previously described (Valjent et al., 2000). After three 10-min washes in Tris-buffered saline (TBS, 0.10 M Tris, 0.14 M NaCl, pH 7.4), free floating brain sections were incubated for 5 min in TBS containing 30 mL.L^−1^ H_2_O_2_ and 10 mL.L^−1^ methanol, and rinsed again in TBS 3 times for 10 min. They were then incubated for 15 min in 2 mL.L^−1^ Triton X-100 in TBS, rinsed 3 times in TBS and blocked in 30 g.L^−1^ bovine serum albumin in TBS. Next, they were incubated overnight at 4°C with primary antibody diluted in the same blocking solution. After three 15-min washes in TBS, they were incubated 1 h with fluorescent secondary antibodies. After three additional washes, sections were mounted in Vectashield (Vector laboratories). Primary antibodies were rabbit Fos antibodies (1:1000, Santa Cruz, #sc-52) and chicken GFP antibodies (1:1000, Thermo Fisher, #A-10262). Secondary antibodies were anti-rabbit A488-labeled antibody (1:400, Invitrogen, #A-21206) and anti-chicken Cy3-labeled antibody (1:800; Jackson Immuno Research, #703-165-155). Images were acquired using a Leica SP5-II confocal microscope.

**Figure 2.**
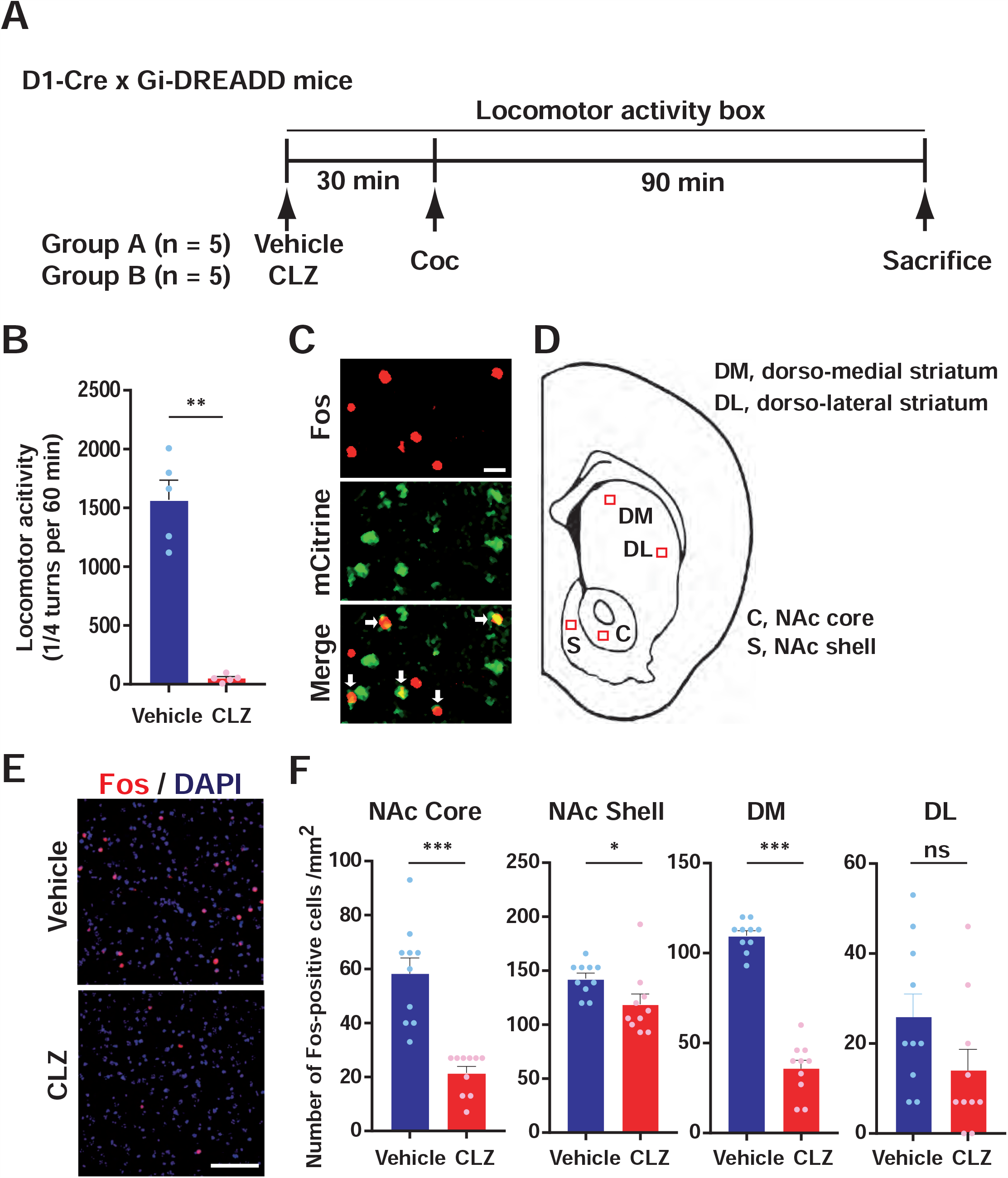
CLZ decreases cocaine-induced Fos expression in Gi-DREADD-expressing striatal neurons. **A)** Schematic experimental design for immunofluorescence. Vehicle or CLZ was injected 30 min before cocaine administration. Mice were killed 90 min after cocaine administration for immunofluorescence. **B**) Inhibition of D1-neurons prevents cocaine-induced hyperlocomotion. Locomotor activity was measured during 60 min after cocaine injection. Data were collected in 5-min bins and shown for each mouse. Bars indicate means + SEM, n = 5 mice per group. Mann-Whitney’s U = 0, *p* = 0.0079, **. **C)** In a mouse injected with vehicle and cocaine, Fos is activated in many cells which express DREARDD under the control of *Drd1* promoter. Representative confocal images in the dorsomedial striatum. Double immunolabeling of Fos (red) and mCitrine (green). Merged image demonstrate colocalization (arrows) of Gi-DREADD-mCitrine and Fos. **D**) c-Fos-positive neurons were stained by immunocytochemistry 90 min after cocaine administration in various areas of the striatum at the 0.98 mm antero-posterior coordinate. The position of the striatal regions used for quantification is indicated by red squares on a mouse brain coronal section (after Paxinos and Franklin, 2001). **E**) Representative images of Fos immunoreactivity (red) and DAPI (blue) in dorsomedial striatum (DM) 90 min after cocaine (20 mg/kg i.p.) administration. **F**) Fos-positive neurons were counted in the 4 regions (388 × 388 µm) indicated in D. Data correspond to 2 regions (right and left) per mouse for 5 mice per group with bar indicate means + SEM. Mann-Whitney’s test, NAc core, U = 0, *p* < 0.0001, NAc shell, U = 17, *p* = 0.01, DM, U = 0, *p* < 0.0001, DL, U = 26, *p* = 0.067. ns, not significant, \**p*<0.05, \*\*\**p*<0.001. Scale bars, C, 20 µm, E, 100 µm.

### Statistical analysis

When normal distribution was not rejected by d’Agostino-Pearson and Shapiro-Wilk tests, data were analyzed with Student’s t test or ANOVA followed by post hoc comparisons as indicated in the figure legends. When normal distribution was rejected, analysis was done with non-parametric tests. All statistical analyses were performed using Prism version 5.00 for Windows (GraphPad Software, La Jolla, CA, USA). A critical value for significance of p < 0.05 was used throughout the study.

## Results

### Gi-DREADD stimulation prevents Fos induction in the dorsal striatum and NAc

We assessed the functionality of Gi-DREADD after a cocaine injection by comparing the number of neurons positive for Fos, a marker of activated neurons, in the striatum of Drd1-Cre x Gi-DREADD mice pretreated with CLZ (0.1 mg/kg) or vehicle. Double transgenic mice were treated with CLZ or vehicle and 30 min later with cocaine (**Fig. 2A**). We observed that cocaine markedly increased locomotion in the group of vehicle-pretreated mice whereas CLZ-pretreated mice displayed little locomotor activity (**Fig. 2B**). In both vehicle- and cocaine-treated mice most Fos-positive neurons, but not all, co-expressed mCitrine, and thus Gi-DREADD, indicating that Fos was mainly induced in D1-neurons as previously reported (Bertran-Gonzalez et al., 2008; Guez-Barber et al., 2011; Zhang et al., 2006, **Fig. 2C**). We counted the number of Fos-positive cells in different regions of the dorsal striatum and NAc (**Fig. 2D**). Compared with vehicle, CLZ pretreatment significantly decreased the number of Fos-positive cells in the NAc core, NAc shell, and dorso-medial striatum (**Fig. 2E, 2F**). In the dorso-lateral striatum, the density of Fos-positive neurons was low and the CLZ-induced decrease did not reach statistical significance. These results indicated that stimulating Gi-DREADD by CLZ inhibited the cocaine-induced activity of D1R-neurons in the dorsal striatum and NAc.

### Inhibition of D1-neurons decreases basal and acute cocaine-induced locomotor activity

Next, we analyzed how inhibition of D1-neurons in Drd1-Cre x Gi-DREADD mice affected spontaneous locomotion and locomotor responses after the first and second cocaine injection (see experimental plan in **Fig. 1B**). On Day 0, during the 30-min period after CLZ pretreatment (Groups 3 and 4) the basal locomotor activity of mice was decreased as compared with vehicle-pretreated mice (Groups 1 and 2, **Fig. 3A, B**). During the 60 min after cocaine injection, locomotor activity was dramatically decreased in CLZ-pretreated mice (Group 4) as compared to vehicle-pretreated mice (Group 2, **Fig. 3A, C**). Importantly, CLZ injection had no effect in wild type animals (WT), which did not express hM4Di: it did not alter basal locomotor activity (**Fig. 3B**, Groups 5 and 6) or cocaine-induced hyper-locomotion (**Fig. 3C**, Group 6). Thus inhibition of hM4Di-expressing D1-neurons by CLZ decreased the spontaneous locomotor activity and acute cocaine-induced hyper-locomotion.

**Figure 3.**
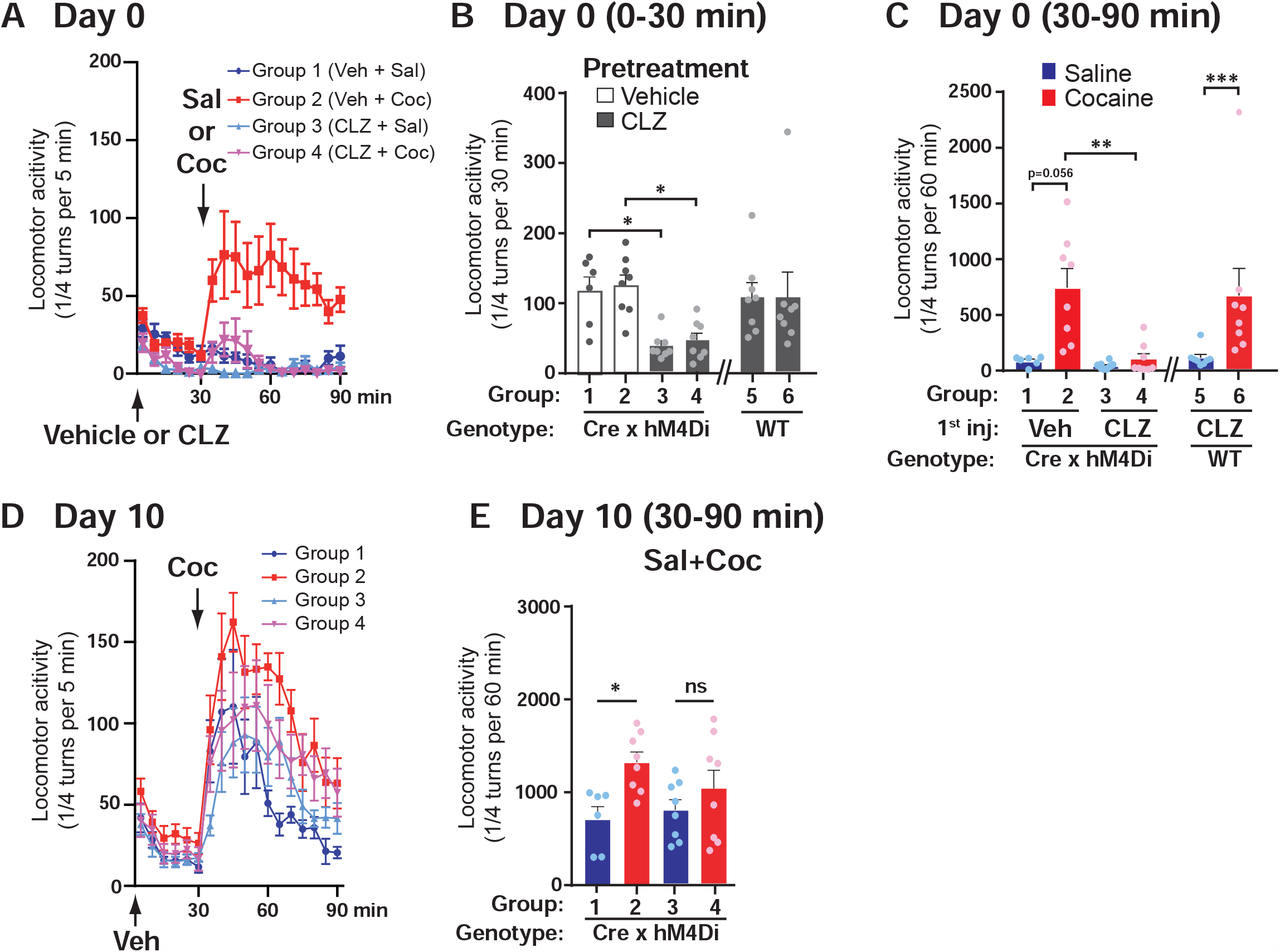
Gi-DREADD-induced inhibition of D1-neurons decreases acute cocaine-induced hyper-locomotion and cocaine sensitization in the TIPS protocol. **A-C)** On day 0, mice were injected with vehicle (groups 1 and 2) or CLZ (0.1 mg/kg, groups 3-6) and placed in the locomotor activity box. Thirty min later, they were injected with saline (groups 1, 3, and 5) or cocaine (20 mg/kg, groups 2, 4, and 6). Group 1, n = 6, groups 2-6, n = 8 mice. (**A**) Time-course of locomotor activity measured in 5-min bins. Repeated measures 2-way ANOVA, interaction, F_(51, 442)_ = 3.43, p < 10^−4^, time, F_(17, 442)_ = 2.91, p < 10^−4^, groups, F_(3, 26)_ = 14.83, p < 10^−4^. **(B**) Total locomotor responses during the 30 min before cocaine injection, one point per mouse, bars indicate means + SEM. Transgenic mice (groups 1-4), Kruskal-Wallis test value = 17.6, p = 5×10^−4^, Dunn’s multiple comparisons test, * p < 0.05. (**C**) Total locomotor responses during the 60 min after cocaine injection (as in B). Transgenic mice (groups 1-4), Kruskal-Wallis test value = 18.4, p = 4 × 10^−4^, Dunn’s multiple comparisons test, ** p < 0.01. Wild type mice (Groups 5, 6, independent experiment) Mann-Whitney test, U = 2, p = 6 × 10^−4^. **D-E**) On day 10, all mice were injected with vehicle, put in the locomotor activity box and then injected with cocaine (20 mg/kg) 30 min later. **D**) Time-course of locomotor activity measured in 5-min bins. Repeated measures 2-way ANOVA, interaction, F_(51, 442)_ = 1.13, p = 0.26, time, F_(17, 442)_ = 25.68, p < 10^−4^, groups, F_(3, 26)_ = 3.97, p = 0.02. **E**) Total locomotor responses during the 60 min after cocaine injection (as in B). Transgenic mice (groups 1-4), Kruskal-Wallis test value = 8.5, p = 0.04, Dunn’s multiple comparisons test, * p < 0.05.

### D1-neurons inhibition impairs the induction of cocaine locomotor sensitization

On Day 10, we injected cocaine to all Drd1-Cre x Gi-DREADD mouse groups to assess the effects of D1-neurons inhibition during the first injection of cocaine on locomotor sensitization, tested by a second injection on Day10 (TIPS protocol, Valjent et al., 2010, **Fig. 3D**). All mice displayed an increased locomotor activity after cocaine injection at Day10 (**Fig. 3D**). In the groups that had not received CLZ (Groups 1 and 2), locomotor response to cocaine on Day10 was significantly increased in the mice that were injected with cocaine for the second time (Group 2) as compared to those that had previously received saline (Group 1, **Fig. 3D, E**). This result confirmed the clear locomotor sensitization with the TIPS protocol in mice (Valjent et al., 2010). In contrast, in the mouse groups pretreated with CLZ on Day 0 (Groups 3 and 4), the responses to cocaine were not significantly different whether or not they had received cocaine a week before. Thus sensitization was blunted in the Group 4 mice that received cocaine on Day 0 in the presence of CLZ (**Fig. 3E**).

### Inhibition of D1-neurons blocks previously sensitized cocaine-induced locomotion

To assess the acute effects of D1-neuron inhibition on locomotor activity in mice previously exposed to cocaine we injected on Day 11 the same mice as in Fig. 3 with CLZ, followed 30 min later by cocaine (**Fig. 4A**). In the four DREADD-expressing groups (1-4), the locomotor responses (30-90 min) were lower than observed the day before in the absence of CLZ (Day 10, compare **Fig. 3 D, E** and **Fig. 4A, B**) and there was no significant difference between groups on Day 11 (**Fig. 4B**). In contrast, in WT mice injected with CLZ (groups 5 and 6), cocaine induced a clear hyper-locomotion (**Fig. 4B**), ruling out a non-specific pharmacological effect of CLZ. Paired analysis showed that locomotor responses to cocaine on Day 11 were significantly decreased compared with those on Day10 in the four Drd1-Cre x Gi-DREADD groups (**Fig. 4C**), including group 2 which was clearly sensitized on Day 10 (**Fig. 3D-E**). These results showed that inhibition of D1-neurons with Gi-DREADD strongly decreased the cocaine-induced hyper-locomotion, including the expression of established cocaine-sensitized locomotion.

**Figure 4.**
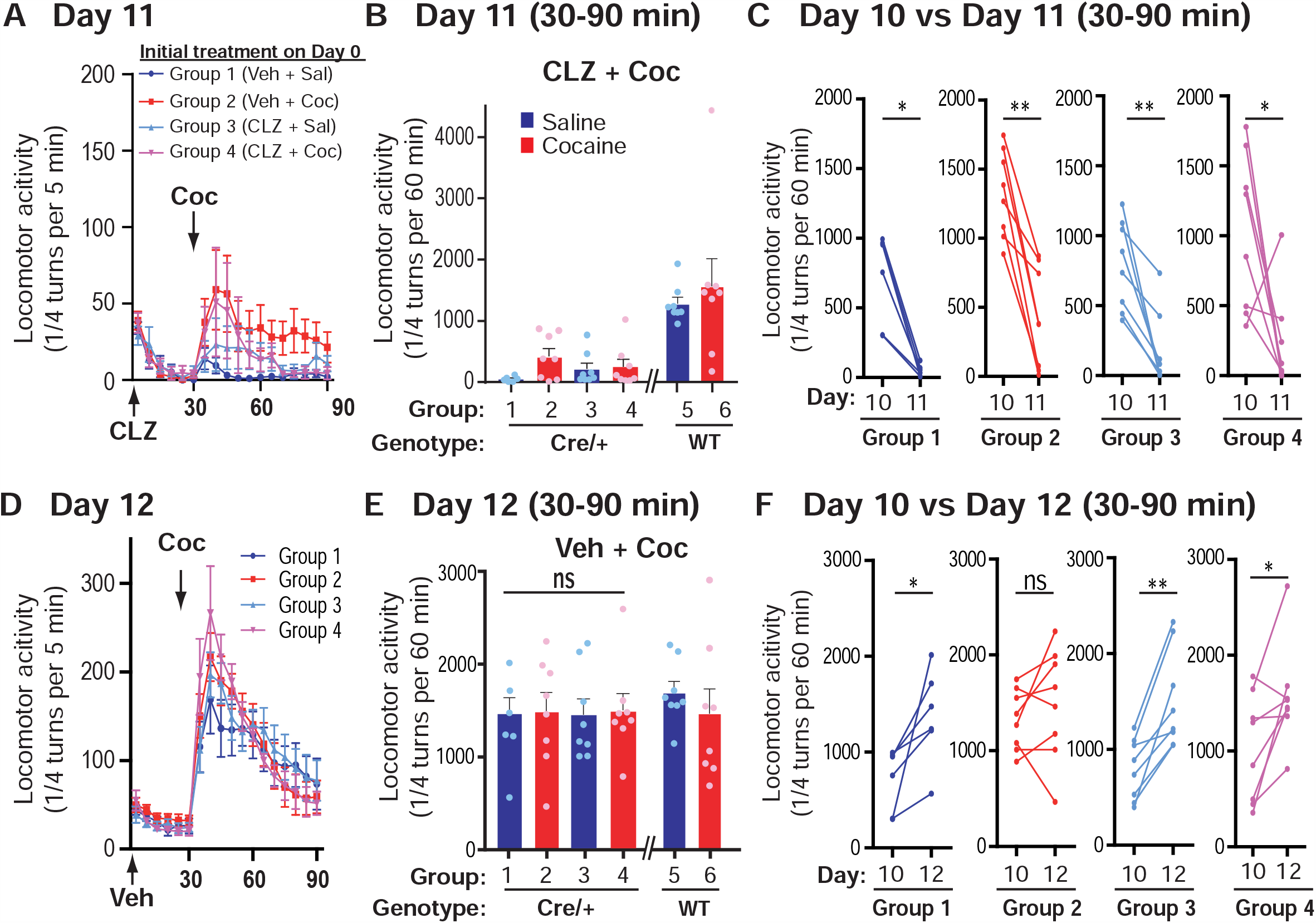
Gi-DREADD-induced inhibition of D1-neurons blocks the expression of previously sensitized cocaine-induced locomotion but does alter its further expression. **A)** On day 11, the same mice as in Fig. 3 were injected with CLZ, placed in the locomotor activity box, and 30 min later, injected with cocaine (20 mg/kg). The time-course of locomotor activity was measured in 5-min bins. Repeated measures 2-way ANOVA, interaction, F_(51, 442)_ = 0.78, p = 0.87, time, F_(17, 442)_ = 4.28, p < 10^−4^, groups, F_(3, 26)_ = 1.82, p = 0.17. **B**) Total locomotor responses during the 60 min after cocaine injection (in A), one point per mouse, bars indicate means + SEM. Transgenic mice (groups 1-4), Kruskal-Wallis test value = 4.2, p = 0.24. Wild type mice (Groups 5, 6, independent experiment) Mann-Whitney test, U = 24, p = 0.44. **C)** Pairwise comparison of each mouse locomotor responses to cocaine on day 10 without CLZ pretreatment (data from Fig. 3E) and on day 11 with CLZ (data from Fig. 4B). Wilcoxon matched-pairs test, Group 1, p = 0.03, Group 2, p = 0.008, Group 3, p = 0.008, Group 4, p = 0.04. **D**) On day 12, the same mice as in A were injected with vehicle, put in the locomotor activity box, and 30 min later, injected with cocaine (20 mg/kg). The time-course of locomotor activity was measured in 5-min bins. Repeated measures 2-way ANOVA, interaction, F_(51, 442)_ = 1.17, p = 0.20, time, F_(17, 442)_ = 44.41, p < 10^−4^, groups, F_(3, 26)_ = 0.13, p = 0.94. **E**) Total locomotor responses during the 60 min after cocaine injection, as in (B). Transgenic mice (groups 1-4), Kruskal-Wallis test value = 2.5, p = 0.78. Wild type mice (Groups 5, 6, independent experiment) Mann-Whitney test, U = 17, p = 0.13. **F**) Pairwise comparison of each mouse locomotor responses to cocaine on day 11 without CLZ pretreatment (data from Fig. 4B) and on day 12 (data from Fig. 4E). Wilcoxon matched-pairs test, Group 1, p = 0.03, Group 2, p = 0.4, Group 3, p = 0.008, Group 4, p = 0.02. Group 1, n = 6, groups 2-6, n = 8 mice. In **E** and **F**, * p < 0.05, **, p < 0.01, n.s. not significant.

### Inhibition of D1-neurons on Day 11 does not alter established cocaine sensitization on Day 12

Cocaine TIPS is context-dependent (Valjent et al., 2010). Context memory retrieval in the presence of a protein synthesis inhibitor can disrupt conditioned locomotor sensitization in rats (Bernardi et al., 2007). We therefore tested whether D1-neuron inhibition on Day 11 by CLZ in Drd1-Cre x Gi-DREADD mice affected the expression of cocaine sensitization the next day (Day 12). On Day 12, we injected all the mouse groups with vehicle and 30 min later with cocaine. There was no significant difference in locomotor activity (30-90 min) between the 4 groups (**Fig. 4D, E**). These results were similar to those observed in WT mice receiving the same treatment on Day 12 (groups 5 and 6, **Fig. 4E**). Paired analysis showed that the locomotor response to cocaine (30-90 min) on Day 12 was significantly increased in Groups 1, 3, and 4, as compared to Day 10 (**Fig. 4F**), indicating the existence of a sensitization step. No significant difference between Day 10 and 12 was observed in group 2 (**Fig. 4F**), suggesting that sensitization had reached a plateau in this group. None of the groups displayed any decreased cocaine-induced locomotor activity on Day 12 compared with that on Day 10. These results showed that the inhibition of D1-neurons with cocaine in cocaine-paired context on Day 11 did not diminish previously established sensitization.

## Discussion

In this study, we used double transgenic mice with conditional expression of Gi-DREADD in neurons that express Drd1 and showed that inhibition of these neurons decreased spontaneous locomotor activity and acute cocaine-induced hyper-locomotion. Inhibition of D1-neurons during the first cocaine administration blunted sensitization to a cocaine challenge ten days later. In addition, acute inhibition of D1-neurons prevented the cocaine-induced locomotor activity of previously sensitized mice. However, inhibition of D1-neurons concomitant with cocaine injection in the drug-associated environment did not diminish previously acquired sensitization. Thus we provide evidence that D1-neuron activity is necessary for acute locomotor effects of cocaine and for induction and expression of psychomotor sensitization.

For the pharmacological activation of Gi-DREADD, we used low doses of CLZ instead of commonly used CNO. CNO induces cellular DREADD-mediated effects with different time courses in the brain and peripheral organs (Alexander et al., 2009). The effects on DREADD-expressing neurons induced by systemic CNO administration appear to result mainly from the action of CLZ produced by the metabolic degradation of CNO (Gomez et al., 2017). CLZ shows a better ability to cross the blood brain barrier and has a higher affinity for DREADDs than CNO (Gomez et al., 2017). Accordingly, in a recent fMRI study of DREADD-induced activation of D1-neurons, we confirmed a more rapid and efficient effect of low doses of CLZ as compared to CNO (Nakamura et al., 2020). We therefore used here low doses of CLZ (0.1 mg/kg, i.p.) to inhibit Gi-DREADD-expressing D1-neurons and observed clear effects of this treatment only in mutant mice expressing Gi-DREADD. Although CLZ has numerous endogenous targets at higher doses including serotonin, dopamine, and muscarinic acetylcholine receptors (Schotte et al., 1993; Bonaventura et al., 2019), in our experiments we did not observe any effect of the low dose of CLZ used in wild type mice, ruling out off target effects.

The role of D1-neurons in cocaine and amphetamine responses has been indirectly studied using *Drd1* knockout mice. In these knockout mice, spontaneous locomotor activity is either unchanged (Drago et al., 1996; Miner et al., 1995) or increased (Xu et al., 1994, 2000; Karlsson et al., 2008) but in most reports the locomotor response to cocaine is lost in these mutant mice (Xu et al., 2000; Karlsson et al., 2008) as well as the ability of cocaine to induce Fos expression in striatal neurons (Drago et al., 1996). *Drd1* knockout mice show reduced sensitized locomotor responses to cocaine (Karlsson et al., 2008) and amphetamine (Karper et al., 2002). These studies explored the role of Drd1 in the action of psychostimulant drugs but not directly that of the neuronal populations which bear these receptors. Recent advances in cell-type-specific technologies allow better characterization of brain regions and cell types involved in specific molecular and functional processes mediating the effects of drugs of abuse, mainly psychostimulants (reviews in Hyman et al., 2006; Durieux et al., 2011; Lobo and Nestler, 2011; Yager et al., 2015). Optogenetic inhibition of striatal D1-but not D2-neurons inhibited the expression of cocaine-induced locomotor sensitization, but its induction was not investigated (Lee et al., 2017). Ferguson et al. injected into the dorsal striatum a viral vector expressing Gi-DREADD under the control of the dynorphin gene promoter and showed that decreasing excitability of striatonigral pathway neurons with CNO did not change spontaneous activity or acute response to amphetamine, but impaired behavioral sensitization (Ferguson et al., 2011). In our study, we observed a strong effect of inhibiting D1-neurons on spontaneous locomotor activity and acute response to cocaine. These stronger effects may result from the wider inhibition of D1-neuron excitability throughout the brain, including in dorsal striatum, NAc and cerebral cortex, in Drd1Cre x Gi-DREADD mice, in contrast to the restricted inhibition in the dorsomedial striatum in the study of Ferguson et al. In addition, it is possible that the efficacy of CLZ to activate DREADD is higher and more rapid than that of CNO, as suggested by Gomez et al. (2017).

Decreasing activity of the medial prefrontal cortex afferent to the NAc did not impair amphetamine sensitization using Cre dependent Gi-DREADD adeno-associated virus and CNO (Kerstetter et al., 2016). In contrast, increasing activity of the striatopallidal neurons (D2-neurons) with Gs-DREADD prevented amphetamine sensitization (Farrell et al., 2013). Combined with our current findings, this report underlines the importance of the balance between D1- and D2-neurons in regulating sensitization. Further studies are needed to determine the regional and cell-type specific roles of D1-neurons in the spontaneous behavior and various behavioral responses to cocaine.

Drug addiction is a complex disorder in which pathological memories termed “addiction memories” play a critical role in the maintenance of addictive behaviors and relapse to drug seeking (reviews in Hyman et al., 2006; Robbins et al., 2008; Tronson and Taylor., 2013). Many rodent studies aimed to erase or weaken these drug-associated memories using pharmacological or non-pharmacological approaches by inducing targeted disruption of memory reconsolidation processes. Memory reconsolidation is the process by which memories become labile under certain conditions of reactivation (retrieval) before being stabilized again and persistently stored in the brain (review in Lee et al., 2017). It was demonstrated that drug-induced conditioned place preference was attenuated or erased by combining re-exposure to the drug-paired stimuli and/or the drug itself with inhibition of protein synthesis (Valjent et al., 2006; Dunbar and Taylor., 2016), ERK signaling (Valjent et al., 2006), D1 receptors (Yan et al., 2014, Marion-Poll et al., 2019) or knockdown of the immediate early gene *zif268/Egr1* (Lee et al., 2005, 2006; Théberge et al., 2010). Locomotor sensitization does not appear to include a clear reconsolidation phase sensitive to protein synthesis inhibitors (Valjent et al., 2006) except when the animals are briefly exposed to the drug-associated context without drug (Bernardi et al. 2007). Here established cocaine-induced locomotor sensitization was not attenuated by a session of re-exposure combined with Gi-DREADD inhibition of D1-neurons. Cocaine locomotor sensitization involves potentiation of excitatory synapses onto nucleus accumbens D1-neurons and can be erased by optogenetic depotentiation of cortical inputs to these neurons (Pascoli et al., 2011). Complete DREADD-induced inhibition of D1 neurons does not mimic this situation.

In conclusion, our results confirm that specific inhibition of D1-neurons decreases spontaneous and cocaine-induced locomotor activity. Moreover, we show that inhibition of D1-neurons alters locomotor sensitization induced by a single injection of cocaine. Our study emphasizes the importance of D1-neurons in the action of cocaine and suggests that selective manipulation of the excitability of these neurons is an interesting therapeutic approach in the context of addiction.

## Acknowledgments

Research in the laboratory of JAG was supported by Inserm and Sorbonne University and by grants from *Fondation pour la Recherche Médicale* (FRM) and ANR (Epitraces). Equipment at the IFM was funded in part by *Fondation pour la recherche sur le cerveau* (FRC) and Rotary *Espoir en Tête*, and by DIM *Cerveau et Pensée Région Ile-de-France*. YN was recipient of a Uehara Memorial Foundation fellowship and a Fyssen Foundation fellowship.

## Competing Interests

The authors declare no competing interest.

## Authors contributions

Yukari N carried out experiments, analyzed results and wrote the manuscript, SL carried out experiments and analyzed results, AN analyzed results, DH, JAG, and Yuki N planned and conceived experiments, analyzed results and wrote the manuscript, Yuki N carried out some experiments

## Abbreviations

CLZ: clozapine
CNO: clozapine-N-oxide
D1-neurons: Dopamine D1 receptor-expressing cells
Drd1: Dopamine D1 receptor
Drd1-Cre mice: *Drd1-Cre* transgenic mice
DREADD: designer receptor exclusively activated by designer drugs
NAc: nucleus accumbens
TIPS: two-injection protocol of sensitization

## Notes

### Competing Interest Statement

The authors have declared no competing interest.

